# PhotoModPlus: A webserver for photosynthetic protein prediction from a genome neighborhood feature

**DOI:** 10.1101/2020.05.10.087635

**Authors:** Apiwat Sangphukieo, Teeraphan Laomettachit, Marasri Ruengjitchatchawalya

## Abstract

Identification of photosynthetic proteins and their functions is essential for understanding and improving photosynthetic efficiency. We present here a new webserver called PhotoModPlus as a platform to predict photosynthetic proteins via genome neighborhood networks (GNN) and a machine learning method. GNN facilitates users to visualize the overview of the conserved neighboring genes from multiple photosynthetic prokaryotic genomes and provides functional guidance to the query input. We also integrated a newly developed machine learning model for predicting photosynthesis-specific functions based on 24 prokaryotic photosynthesis-related GO terms, namely PhotoModGO, into the webserver. The new model was developed using a multi-label classification approach and genome neighborhood features. The performance of the new model was up to 0.872 of F1 measure, which was better than the sequence-based approaches evaluated by nested five-fold cross-validation. Finally, we demonstrated the applications of the webserver and the new model in the identification of novel photosynthetic proteins. The server was user-friendly designed and compatible with all devices and available at http://bicep.kmutt.ac.th/photomod or http://bicep2.kmutt.ac.th/photomod.

## Introduction

Photosynthesis is a complex and diverse process developed in green organisms for converting sunlight into useable chemical form. Discovering new components in the photosynthetic process is substantially important, as it may point the way to improve photosynthesis efficiency [1-4] and its applications [5-8].

Considering the efficient pipeline used in the identification of novel photosynthetic proteins, it is clear that computational approaches are crucial as a first screening step before a more in-depth study with experimental approaches [9-11]. An earlier attempt of the computational approach has been made based on sequence similarity search. It is trustable when high sequence similarity is found in the existing database, but its accuracy drops quickly when the lower similarity is found. It was shown that the classification of photosynthetic proteins suffers from the diversity of photosystem, which can cause a high false-positive rate of up to 70% [12]. Moreover, it is inapplicable when no similar sequence is found in the database. Therefore, techniques using different sources of the feature have been developed and shown to overcome the classical sequence similarity-based method [13]. SCMPSP [14] is a genetic algorithm model that uses dipeptide property and amino acid composition to classify photosynthetic proteins. Although SCMPSP performed better than the sequence similarity search method, it was not significantly better than other machine learning models indicating the limitation of the sequence-based feature. Therefore, a new model employing a genome neighborhood network (GNN) as a feature called PhotoMod was developed [15]. PhotoMod combined protein clustering, genome neighborhood conservation scoring, and random forest (RF) algorithm to classify photosynthetic proteins. It was shown that the genome neighborhood-based model outperformed sequence-based approaches and showed the potential to predict novel photosynthetic proteins. However, these methods can only classify proteins into two categories i.e. photosynthetic and nonphotosynthetic protein remaining a big task to investigate functional subclasses in photosynthesis in a laboratory.

It has long been known that photosynthesis consists of several subsystems including photosystem I, photosystem II, electron transport system, ATP synthase, NADH dehydrogenase, light-harvesting complex, carbondioxide fixation and many of assembly factors and regulators. However, many proteins in these systems, for example, proteins in ATP synthase and NADH dehydrogenase, can be homologously observed in nonphotosynthetic organisms. Thus, to systematically define the group of photosynthetic proteins, Ashkenazi S, et al. [12] created the list of functional terms that are unique to photosynthesis based on gene ontology system to identify photosynthetic proteins. The list contains 61 GO terms, which are the descendants of photosynthesis (GO:0015979) and not be the part of other functions that are not photosynthesis or its descendants. Among these 61 GO terms, only 24 GO terms are found in photosynthetic prokaryotes.

The general existing protein function prediction methods may be applied to this particular task, although they were not specifically developed to predict photosynthesis subclasses. For example, SVMprot model uses a support vector machine (SVM) with physicochemical features, such as amino acid composition, hydrophobicity, polarity, polarizability, and charge, of protein sequences to predict 192 functional classes including only three photosynthesis-specific subclasses [16, 17]. DeepGOPlus combines sequence similarity-based predictions with a deep convolutional neural network for large-scale protein function prediction, including 5,210 functional classes and ten photosynthesis-specific subclasses [18]. Nevertheless, these models are less specific to photosynthetic functions covering up to only ten of the total 61 photosynthesis-specific classes [12]. Most importantly, sequence-based models tend to perform worse in biological process (BP), which is the main category (in gene ontology resource) of photosynthesis subclasses [19, 20]. It was also suggested that ensemble methods combining data from different sources could improve prediction accuracy [20]. Therefore, we hypothesize that the combination of data from sequence features and genome neighborhoods can improve prediction accuracy.

In this study, we created a new model called PhotoModGO utilizing multi-label learning and genome neighborhood profile as a feature to predict 24 subfunctional classes of photosynthetic function. We demonstrated that our new model overcomes two sequence-based methods, BLAST and DeepGOPlus, investigated by nested cross-validation. In order to make our model practically applicable, we developed a new web server, namely PhotoModPlus, to serve for photosynthetic protein function identification. The webserver contains two main applications, i) machine learning prediction for predicting photosynthetic function, and ii) GNN generation for visualizing genome neighborhood network (GNN), which allows users to investigate gene neighborhood conservation and infer photosynthetic protein function from the network analysis. PhotoModPlus provides an easy-to-use web interface for input submission and convenient interpretable output to evaluate the potential of proteins in photosynthetic function.

## Method

### Dataset collection

#### Training and test datasets

The photosynthetic protein dataset was retrieved from the UniprotKB database. 15,191 protein sequences consisting of at least one of 61 photosynthesis-specific GO terms identified by Ashkenazi et al. [12] were included in the dataset. To avoid incomplete gene neighborhood identification from incomplete genomes, only photosynthetic proteins from 154 photosynthetic prokaryote complete genomes were included in the dataset, as reported in [15]. To reduce sequence redundancy, we used method USEARCH [21] to cluster those similar sequences (sequence identity ≤ 50% as a diverse dataset and ≤ 70% as an easy dataset). The USEARCH command is

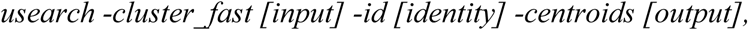

where input is a file in Fasta format, identity is percent sequence identity cutoff to the cluster, and output is representative selected sequences in Fasta format. The number of sequences in the dataset was reduced to 369 sequences with identity ≤ 50% and 1,021 sequences with identity ≤ 70%, which were applied for model development and model comparison.

#### Novel photosynthetic protein dataset

We also collected a set of novel proteins from literature to evaluate model performance. We would consider those proteins as a novel if they had never been collected in the Uniprot database released in September 2016. Since the development version of our model has been trained from the data retrieved from the Uniprot database released in September 2016, we assume that genes annotated after this time are novel. BLAST was used to check their availability in the database with the criteria of sequence identity ≥ 50% and query and subject coverage ≥ 70%. The lack of sequence homolog makes them the ideal for testing the feasibility of our models for facilitating the functional annotation of novel proteins.

### PhotoModGO development

As shown in the previous study [15], we developed a binary classification model, which combines protein clustering, genome neighborhood extracting and scoring, and random forest algorithm to classify photosynthetic proteins. In this study, we developed a new model using a multi-label classification approach, which allows each data point to be assigned to more than one functional class at the same time, to classify photosynthesis subclasses based on a genome neighborhood feature.

The process starts by retrieving photosynthetic protein dataset with photosynthesis-specific GO terms and removing redundant data as described in the dataset collection (Fig 1). Then, the loci of those photosynthetic protein-coding genes were determined and used for calling gene neighborhood. The genes were considered neighbors if they are within 250 bp on the same strand. The two gene clusters can be merged into the same neighborhood gene cluster if they are in a range of 200-1000 bp in the divergent direction following the operon interaction concept, as demonstrated in S1 Fig. The homologous relationship between protein sequences was determined by the protein clustering method with three stringency criteria, according to the previous study (1E-10, 1E-50, and 1E-100)[15]. The genome neighborhood conservation scores (Phylo score) were calculated based on the phylogenetic tree of organisms that conserve those gene neighborhoods (as described in [15]). The quantile cut-off points were determined and used for converting the raw Phylo scores to simple numeric form (i.e., 0, 1, 2, and 3). Hereafter, we call the list of gene neighbors and their Phylo score of the query gene as a genome neighborhood profile. The matrix of the query genes and their genome neighborhood profiles was built and concatenated with the binary matrix of GO term annotations coresponding to the query genes. This processed matrix was converted to ARFF format, which is a standard format for machine learning softwares. Note that we keep using a matrix form to display the data in Fig 1 as it is easier to understand.

**Fig 1.**
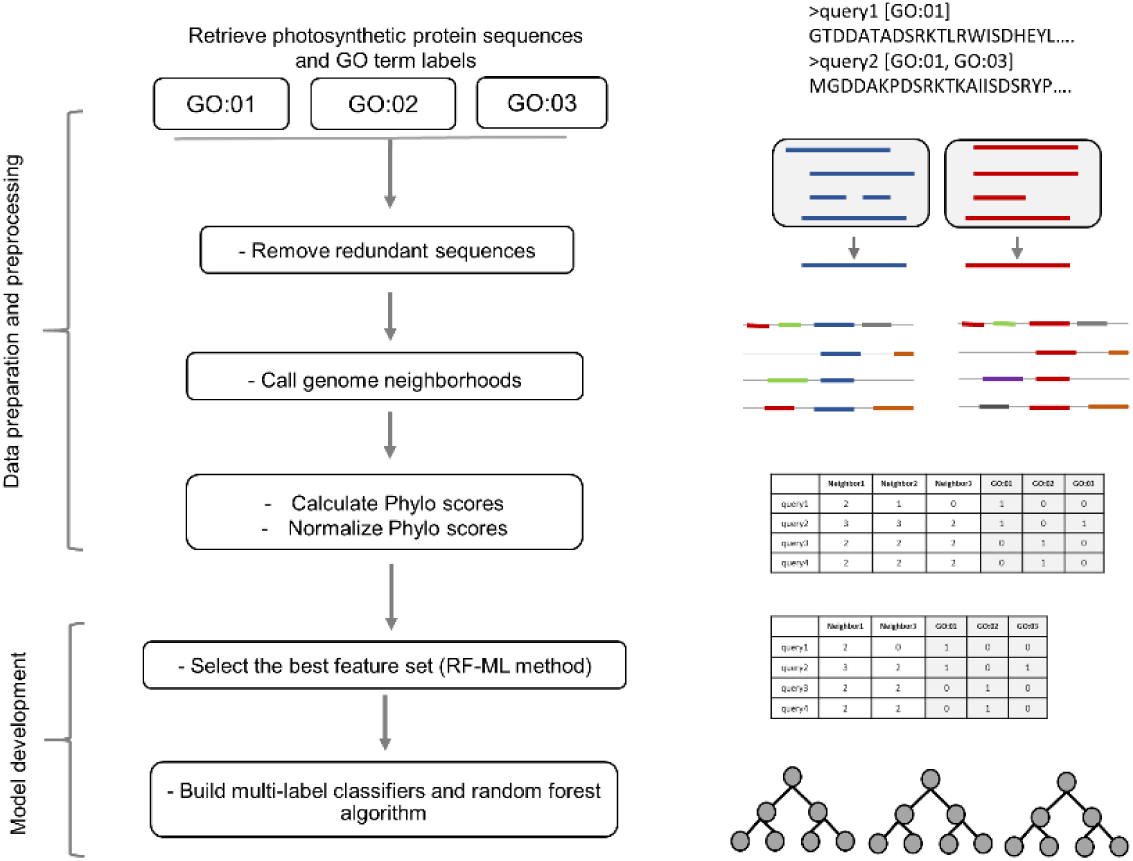
Overview of PhotoModGO development. The process starts by retrieving photosynthetic protein dataset with photosynthesis-specific GO terms and removing redundant data. The loci of those photosynthetic protein-coding genes were determined and used for calling gene neighborhoods. The Phylo scores of the neighboring genes were calculated and then normalized. The genome neighborhood profile containing a list of neighboring genes and their Phylo scores for each query was created and was concatenated as a matrix form. The multi-label feature selection method, RF-ML, was employed before building a multi-label classification model based on a random forest algorithm. The right panel shows the graphical representation for each step.

The next steps are the process for building a machine learning model, which consists of i) selecting the optimal feature set, and ii) training the model. Due to a large number of features in the genome neighborhood profile, we removed redundant and irrelevant features in the first step to increase learning speed and model performance. We employed the multi-label feature selection method, RF-ML [22], which is the extended version of the single-label feature selection method, Relief. The RF-ML method was developed by using the dissimilarity function between multi-labels to determine the optimal set of features. That is, it considers the relationship between labels making the prediction performance of the model better. We modified the original RF-ML script to obtain the top-N features ranked by dissimilarity score, where N is a number of features in the feature subset. The N number was fine-tuned during the model development process.

The next step is the building of a machine learning model, in which we employed a problem transformation method to transform a multi-label dataset into the multi-class dataset allowing usual learning algorithms to learn. We initially selected three different multi-label transformation methods for model development, and the method with the best performance was selected as the final model. The first method is Binary Relevance (BR) [23], a classical approach in multi-label classification. It constructs a binary classifier for each label, and the final prediction is determined by aggregating the classification results from all classifiers. The problem of this method is that it ignores label relationships resulting in poor performance when applied on real-world datasets. The second method is Label Power-set (LP) [24], which it is developed to take label dependency into account by constructing binary classifiers from all possible sets of those labels. Therefore, it usually suffers from worst-case time complexity when the size of the label set is large. The last method is RAndom k labEL sets (RAkEL) [25], an ensemble of LP classifiers, each of which is built from a different random label subset of a size k. Therefore, the classification performance of this method depends on the random strategy of the label subset. We applied these transformation methods with the random forest algorithm as a base binary classifier, which performs the best in our previous study [15]. We developed classification models by using the MEKA framework [26]. Three parameters are tuned to optimize the model, i.e., number of the tree, the number of random features used in the tree in the random forest, and the number of selected features from the RF-ML method.

### Nested cross-validation

We used five repetitions of nested 5×3-fold cross-validation (CV) to evaluate the model performance with the optimal parameter set. The schema of the nested fold CV is demonstrated in S2 Fig. In every fold, the training set is separated to perform nested fold CV to find the best parameter set before validating by the test set. That means we can test the model in each fold with the best parameter set without the leaking of the information during the step of model training.

### Baseline comparison methods

#### DeepGOPlus

DeepGOPlus is a new sequence-based method that uses a deep convolutional neural network algorithm to predict multiple protein functions. The DeepGOPlus was retrieved from https://github.com/bio-ontology-research-group/deepgoplus.git. We newly trained this model by using our photosynthetic protein dataset. Due to the high computational requirement of DeepGOPlus, we selected two optimal parameter sets, fine-tuned from the original paper, to use in this study.

#### BLAST

We also use a sequence similarity method based on Diamond BLAST as a baseline method for comparison. Training data is used as a database, and the test set is used as a query. Annotation from the training dataset is transferred to a similar sequence in the test set if it passes E-value criteria, which range from 20 to 1E-30. The percent sequence identity is used as a probability score. The diamond BLAST command is shown below.

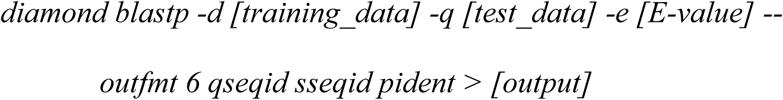

### Evaluation metric

To evaluate prediction performance, we use simple and easily interpretable metrics, F1_max_ [27], which has been recommended by the Critical Assessment of Functional Annotation (CAFA) [28]. F1 score is the harmonic mean between precision and recall, and F1_max_ is the maximum F1 score overall classification thresholds of each model.

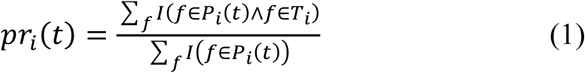

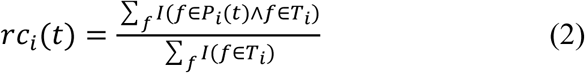

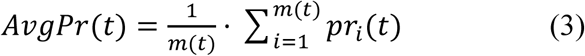

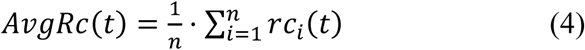

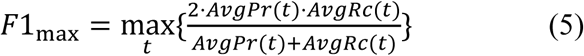

First, precision (*pr*_*i*_) and recall (*rc*_*i*_) are calculated as equations (1) and (2). For every protein *i* in the dataset, *f* is a GO term label, *T*_*i*_ is a set of true GO term labels and *P*_*i*_*(t)* is a set of predicted GO term labels of threshold *t. I* is an identity function which returns 1 for true and 0 for the false condition. Second, average precision (*AvgPr*) and average recall (*AvgRc*) are calculated as equations (3) and (4), where *m(t)* is a number of proteins that contain at least one predicted label and *n* is a total number of proteins. The F1_max_ is calculated as equation (5) with the thresholds *t* in range of 0 to 1 and a step size of 0.1.

### Web implementation

The PhotoModPlus web server was developed using Flask framework version 1.0.2 and python script and implemented using Apache HTTP web server version 2.4.18. PhotoModPlus web server is available at bicep.kmutt.ac.th/photomod and bicep2.kmutt.ac.th/photomod.

## Results

### Performance of PhotoModGO in multi-label classification

In this study, we developed a new multi-label classifier, namely PhotoModGO, to predict functional subclasses in the photosynthetic system. Photosynthetic subclasses were selected from the list of 61 photosynthetic specific GO terms [12], in which only 24 GO terms (S3 Table) belong to prokaryotes. Data preprocessing was carried out as described in the method section.

Among the multi-label classifiers, RAkEL performed the best with the F1_max_ value of 0.872 following by the BR method with a value of 0.862 (Table 1). The performance between these two algorithms was not significantly different (*p*-value = 0.166) observed by the Wilcoxon signed-rank test (S4 Table). LP performed the worst with an F1_max_ value of 0.693, which was significantly different from the two methods (*p*-value < 0.00001). Although the recall value of the LP model was highest, the precision value was lowest compared to the others. It is consistent with the previous benchmark [29] showing that the ensemble transformation methods generally perform better than simple transformation methods.

**Table 1.** Performance comparison of PhotoModGO to other sequence-based models using five replications of nested 5×3 fold-cross validation.

| Model | F1<sub>max</sub> (±SD) | Precision (±SD) | Recall (±SD) |
| --- | --- | --- | --- |
| PhotoModGO-RAkEL | 0.872 (0.035) | 0.926 (0.031) | 0.869 (0.042) |
| PhotoModGO-BR | 0.862 (0.029) | 0.902 (0.037) | 0.873 (0.042) |
| PhotoModGO-LP | 0.693 (0.046) | 0.665 (0.061) | 0.940 (0.021) |
| DeepGoPlus | 0.702 (0.032) | 0.771 (0.041) | 0.693 (0.048) |
| BLAST | 0.603 (0.049) | 0.601 (0.052) | 0.624 (0.050) |

The same dataset was used to train two sequence-based models, DeepGoPlus and BLAST, for performance comparison. We found that DeepGoPlus was able to achieve 0.702 of F1_max_, 0.771 of precision, and 0.693 of recall. The classical BLAST method was the worst-performing method in this comparison with F1_max_, Precision, and Recall of 0.603, 0.601, and 0.624, respectively. Although the result showed that RAkEL and BR methods performed better than DeepGoPlus significantly (*p* < 0.00001), the performance of the LP method was not statistically different from the performance of DeepGoPlus (*p* = 0.135). It is clear that the genome neighborhood-based models outperformed the sequence-based models on average.

### Sequence identity-independent performance of PhotoModGO

The more similar protein sequence tend to have more similar functions [30]. Therefore, we examine the role of sequence identity in our model performance. We created a new dataset containing sequence identity ≤ 70%, called easy dataset. PhotoModGO-RAkEL, DeepGOPlus, and BLAST models were trained with this dataset and evaluated by using five replications of nested 5×3 fold CV. We observed that DeepGoPlus and BLAST methods performed better on the easy dataset (identity ≤ 70%) comparing to identity ≤ 50% dataset, but we cannot observe the difference in PhotoModGO performance (Fig 2), indicating that PhotmoModGO does not suffer from the situation where proteins with similar functions share low sequence identity scores.

**Fig 2.**
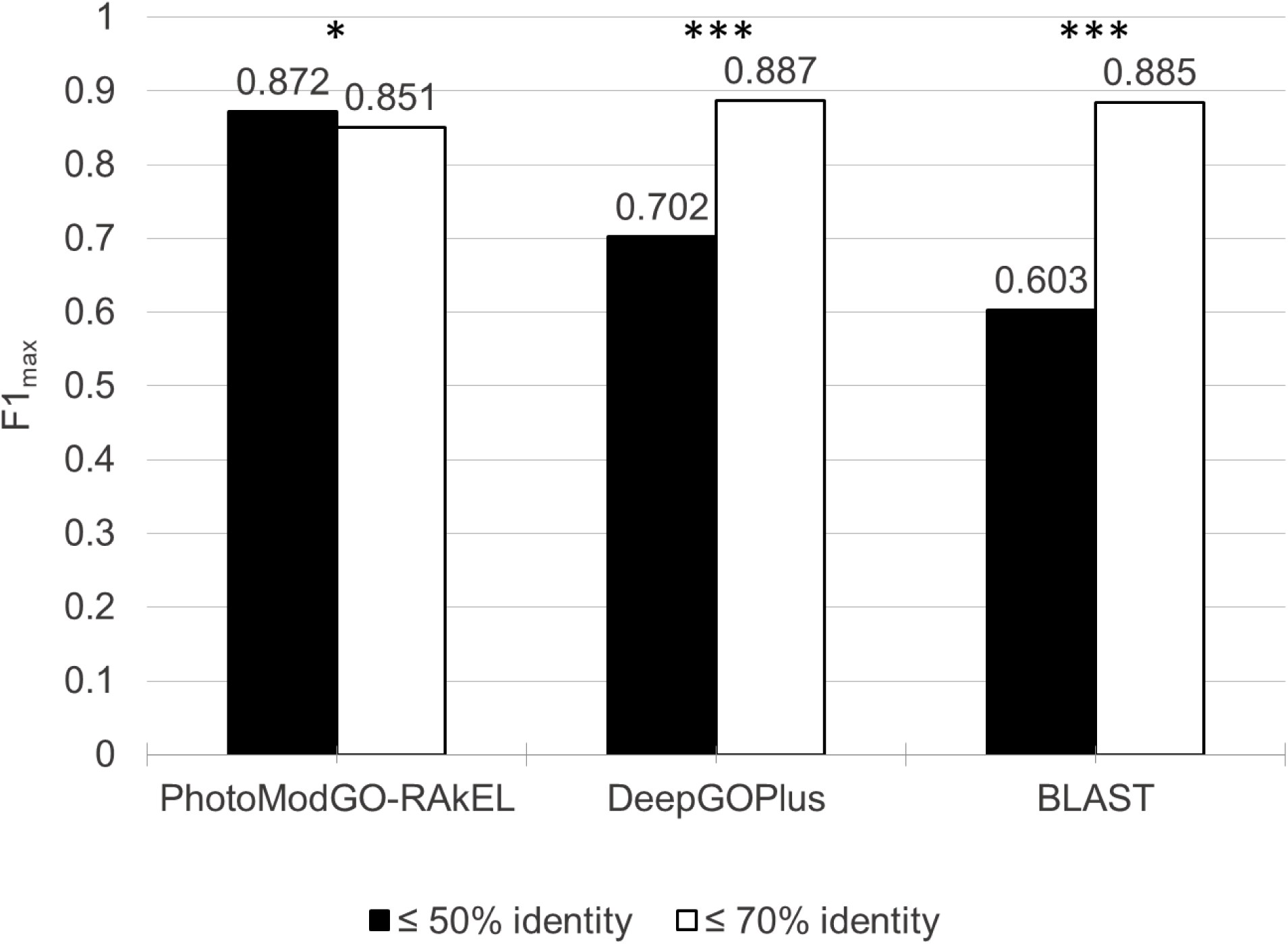
Multi-label classification performance of photosynthesis functions via 5 replications of nested 5×3 fold-cross validation. The unique sequence dataset at level ≤ 50% identity (diverse dataset) is represented by a black bar, while the dataset at level ≤ 70% identity (easy dataset) is represented by a white bar. Asterisks indicate statistically significant difference tested by Wilcoxon signed-rank test. Significant *p*-value are marked with asterisks (*,*0.01< p <*0.05; ***, *p*<0.00001).

DeepGoPlus achieved 0.887 of F1 measure in the easy dataset, while BLAST method can achieve 0.885. This better performance of sequence-based models indicates that they take advantage of the more similar sequence to predict function. On the other hand, PhotoModGO cannot take advantage of such a sequence property resulting in no significant difference between high and low sequence identity datasets in the prediction performance. The reason for this is that PhotoModGO considers both sequence relationships and genome neighborhood profiles for the classification. For example, weakly homologous sequences holding the same genome neighborhood profile are generally classified under the same functional class.

### Functional prediction of novel photosynthetic proteins

We demonstrated the use of our new model by predicting the function of novel photosynthetic genes/proteins. We retrieved 17 recently identified photosynthetic proteins, which have never been collected in the Uniprot database (Sep 2016). As shown in Table 2, the functional annotations of many proteins from literature are crudely described, which makes it difficult to do a further experiment. We can apply the new model to these proteins to specify functional classes related to photosynthesis. We found that many proteins were predicted to be related to PSII, such as RfpA, IflA and DpxA, while the only IsiX was predicted to be related to PSI. Interestingly, sll0272 product has been documented as a homolog of NdhV in *Arabidopsis thaliana*. Deletion of this gene reduced the NADPH dehydrogenase (NDH-1)-dependent cyclic electron transport around photosystem I (NDH-CET) activity, which has previously shown as a protective role against several stress conditions e.g. high light, high salt and high temperature [31-33]. It has been proposed that sll0272 product is a ferredoxin-binding domain of cyanobacterial NDH-1 localized in the thylakoid membrane [31]. Based on our prediction, we propose that the product of this gene functions in the photosynthetic system as a regulator or stabilizer for PSII components. A few studies supporting this prediction showed that the absence of NDH-CET activity impairs PSII repairing process under heat stress conditions [34, 35]. Besides, three proteins i.e. ApcD4, MpeZ and all4940 were predicted to be related to phycobilisome consistent with the function from the literature.

**Table 2.**
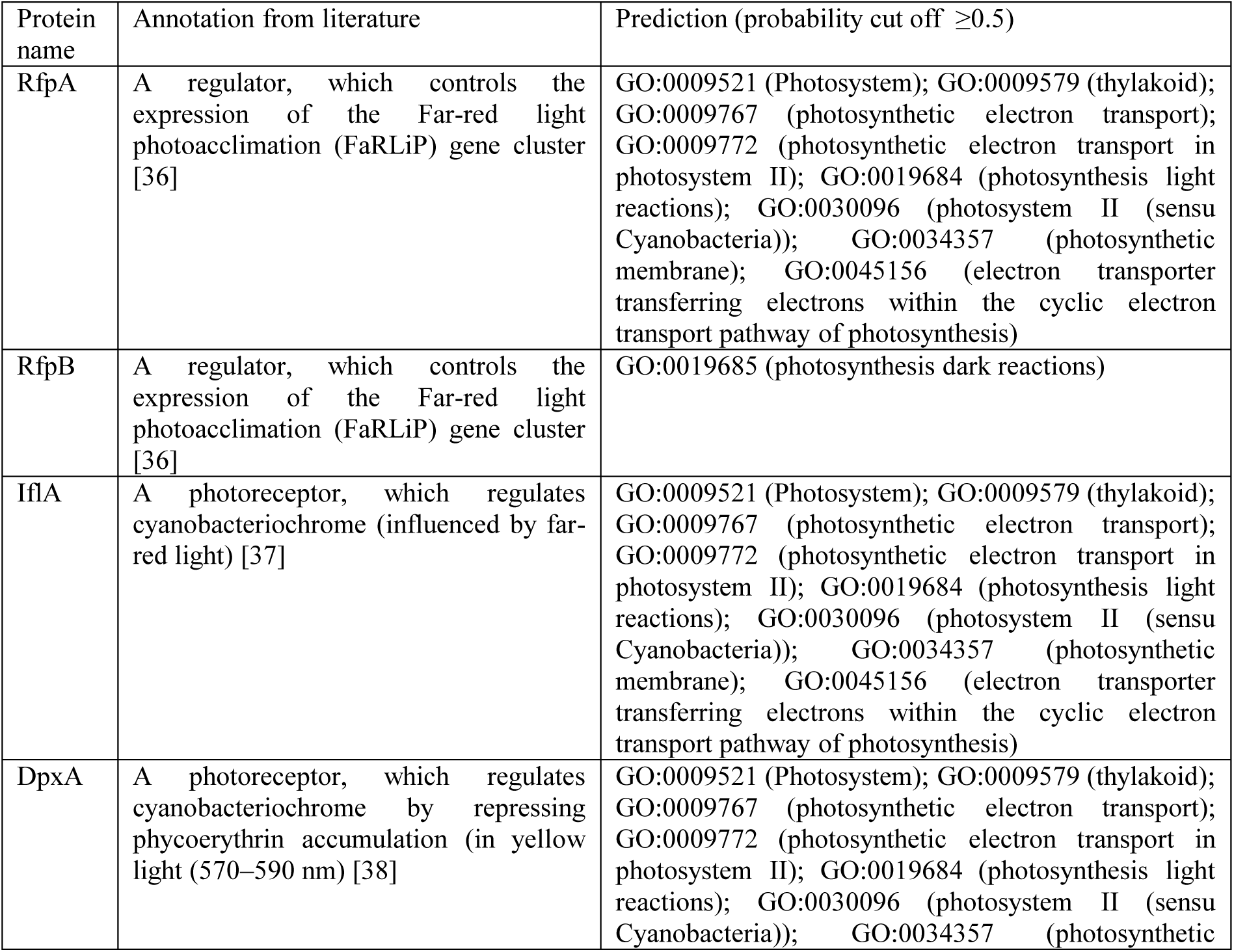
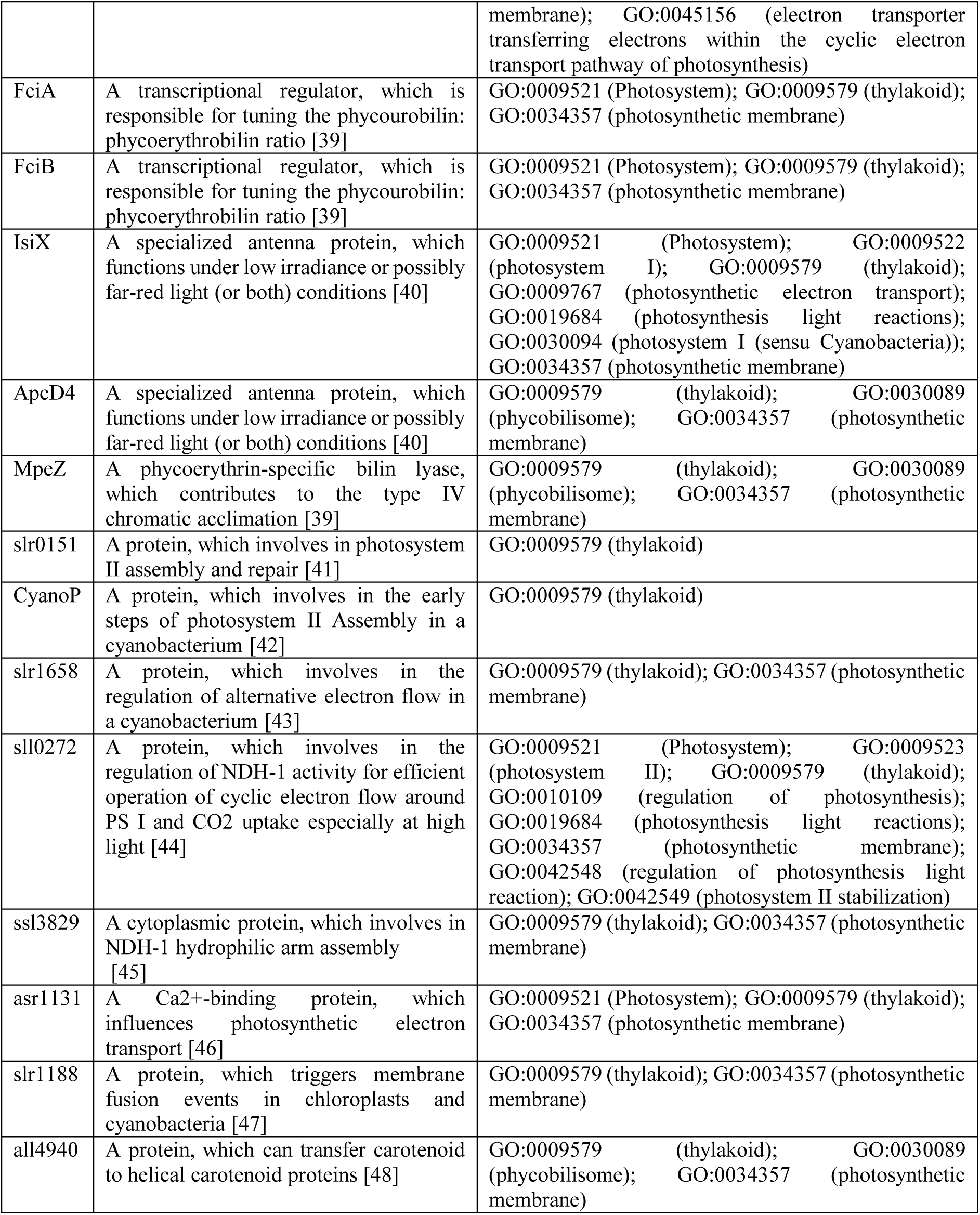
Functional prediction of novel photosynthetic proteins using PhotoModGO.

Surprisingly, we observed that the predicted GO terms from the DeepGoPlus model were sparse and not informative (S5 Table). For example, all of the query genes were assigned by GO:0034357 (photosynthetic membrane) and GO:0009579 (thylakoid), while only ssl3829 and asr1131 were assigned by a more specific term, GO:0009523 (photosystem II). Moreover, only six from 17 query sequences could be assigned with photosynthetic GO terms by SVMprot webserver (S5 Table). These results indicated the advantage of PhotoModGO in the classification of photosynthetic subclasses over the sequence-based approaches.

### Functional prediction of unknown proteins in *Synechocystis* sp. PCC 6803

Identification of novel photosynthetic proteins might provide us a new route to improve photosynthesis efficiency. It has been found that more than 50% (1,885 from 3672) of protein-coding genes in the model organism of photosynthetic prokaryote, *Synechocystis* sp. PCC 6803, are unknown (data from http://genome.microbedb.jp/cyanobase, Sep 2018). Therefore, we applied two machine learning models, PhotoMod and PhotoModGO, implemented in PhotoModPlus webserver to screen the unknown genes in *Synechocystis* sp. PCC 6803 genome to find the potential novel photosynthetic genes. As a result, we could predict ∼100 high confident photosynthetic gene candidates and their function (as shown in S6 Table), some of which are shown in Table 3 as interesting candidates with informative annotations.

**Table 3.**
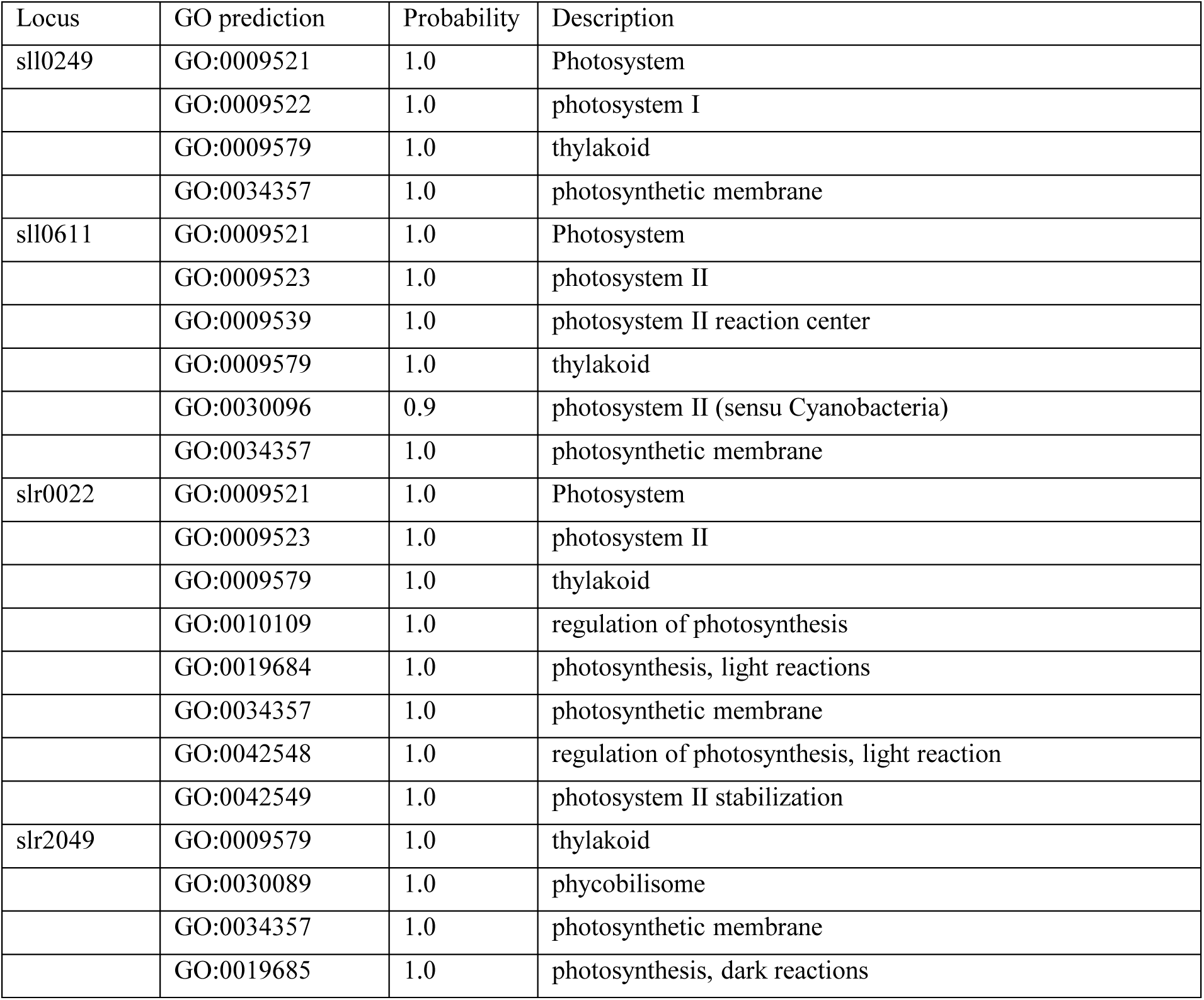
Identification of potential photosynthetic genes in Synechocystis sp. PCC 6803 genome using PhotoModPlus Prediction.

sll0249 localizes next to *isiAB* operon, thus it was designated to *isiC*. While IsiA protein was shown to associate with PSI to form a ring-like complex around a PSI reaction center under iron deficiency condition [49], *isiC* was observed to be transcriptionally enhanced during iron deficiency with no functional designation [50]. Also, the *isiA* mutant was previously reported to be sensitive to high light [51], while the *isiC* mutant showed high sensitivity to oxidative stress [52]. It is known that the oxygenic photosynthetic bacteria consume a large amount of iron for maintaining photosynthesis (e.g. a part of several iron-sulfur cluster-containing proteins). Thus, to minimize harmful radicals from oxidative interaction, iron accumulation in cells should be tightly regulated [53]. According to our prediction result and such pieces of evidence, we proposed that sll0249 or IsiC appears to play an important role in balancing PSI activity and oxidative stress response.

We also predicted that sll0611 functions in PSII. DNA microarrays of *Synechocystis* PCC6803 showed that sll0611 gene responses to the depletion of *lexA* gene, which is the key regulator of the SOS response induced by DNA damage [54]. A more comprehensive study achieved by RNA-seq analysis showed that the depletion of *lexA* gene resulted in altered expression of many genes, including those in photosynthesis, although no significant change of sll0611 was observed in this study [55]. A further study focusing on sll0611 is required to clarify photosynthesis-related function. slr0022 product was previously identified as core cyanobacterial clusters of orthologous groups of proteins (CyOGs) [11] and was predicted by the sequence similarity-based approach as Fe-S cluster protein. Our model was able to predict more specific functions of slr0022 related to photosynthesis. Additionally, our prediction result of slr2049 is consistent with the recent publication showing that slr2049 product functions as lyase to catalyze phycobilin chromophores, which is a part of phycobiliproteins in phycobilisome [56].

### PhotoModPlus web server demonstrations

To increase accessibility, we built PhotoModPlus webserver, which provides two main applications: i) PhotoModPlus prediction providing a classification platform of photosynthesis-related proteins and ii) PhotoModPlus GNN providing efficient GNN visualization.

#### Photosynthetic function prediction by the machine learning model

Our prediction tool provides users to predict photosynthetic proteins by only one click. The prediction pipeline contains three modules: i) binary model to classify photosynthetic protein (PhotoMod), ii) multi-label model to predict photosynthetic subclasses (PhotoModGO) and iii) Interproscan to predict protein function via protein motif search. The single or multiple unknown protein sequences in FASTA format can be used directly as input for the PhotoModPlus web server, where the homologs of the input are identified, and the genome neighborhood profiles are automatically generated. E-value cutoff and the maximum number of BLAST matches are available to adjust for limiting the number of the outputs. Fig 3 illustrates an example of the prediction of an sll0272 protein sequence, which was classified as photosynthetic protein shown with its predicted photosynthesis-specific GO terms.

**Fig 3.**
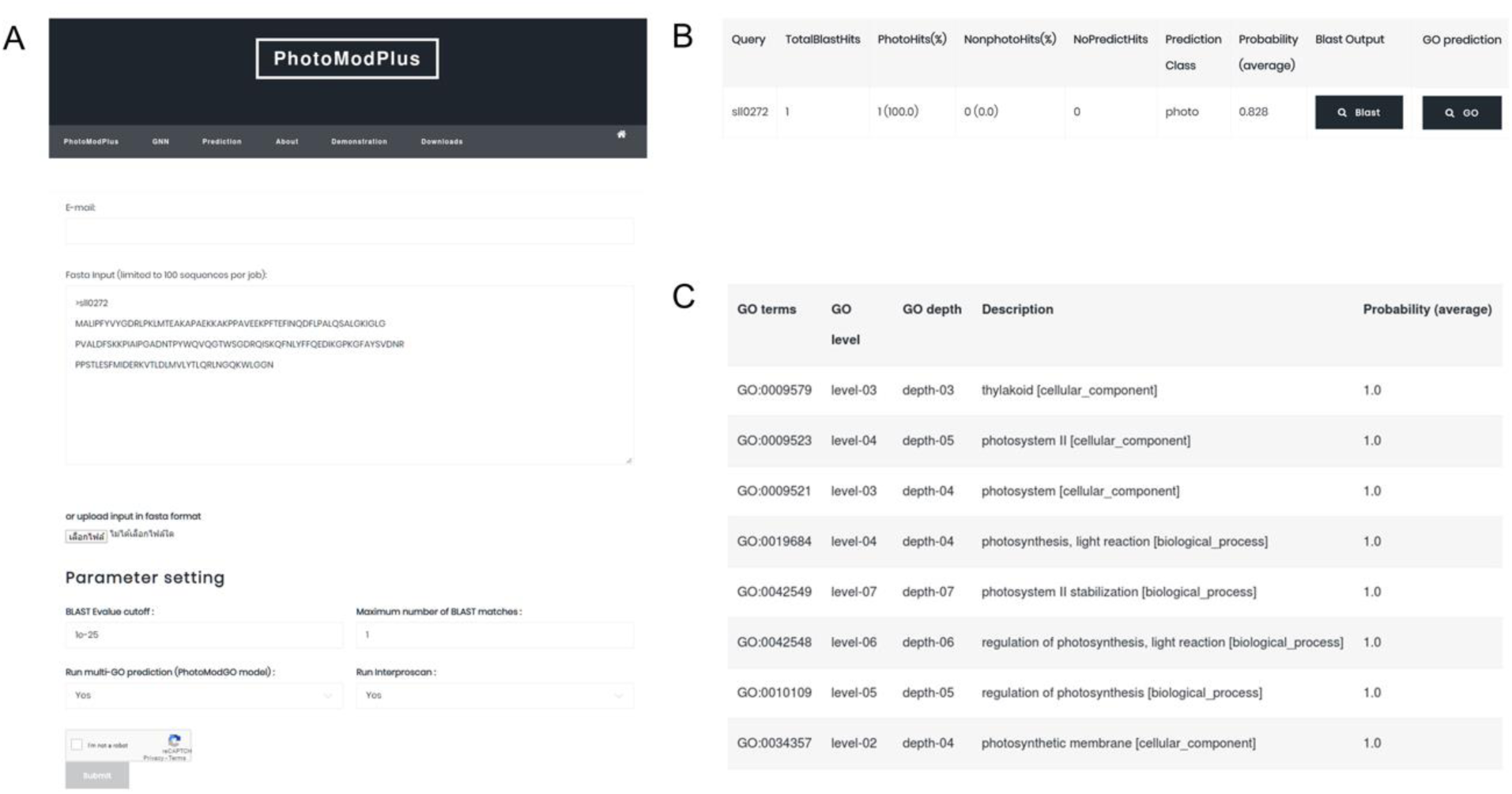
Example of the machine learning prediction output from PhotoModPlus web server of sll0272. (A) illustrates the submission page of the PhotoModPlus web server. Users can submit the input sequence in Fasta format and modify BLAST parameters for matching the query to the sequences in the database. (B) shows the prediction output from the binary classification model (PhotoMod) indicates that sll0272 has a high chance (83%) to be photosynthetic protein. (C) shows the prediction output from the multi-label classification model (PhotoModGO), which indicates the functions of sll0272 in relation to photosystem II and regulation of photosynthesis.

#### Functional guidance by a genome neighborhood network

GNNs were previously shown to facilitate protein function prediction on a large scale and enable the discovery of novel protein function [57-59]. Users can visualize the GNN of interested genes by flexible build-in network visualizer in PhotoModPlus GNN. It accepts single and multiple sequences in familiar FASTA format as input. Users can use default parameters to submit query sequences, and the link to the output page will be sent to the user by email. The summited sequences are blasted to our protein sequence database collected from only photosynthetic organisms. The output is visualized as GNN via Cytoscape.js [60] plugin. The network was designed to be portable, informative, and easy to use. As demonstrated in Fig 4A, the hexagonal node represents the query, while the circular node represents its genome neighbors. The labeled text in each node represents a protein cluster ID. The size of the genome neighbor node varies according to its conservation score (Phylo score). There are three build-in protein clustering cutoffs (E-value: 1E-10, 1E-50, and 1E-100) available on the top of the network to define the level of homologous relationship of proteins. These pre-calculations allow the user to immediately adjust the network according to the stringency. Users can click on the node to observe the most enriched functions in each protein cluster in the network. We also provide a link from the node to the hub of protein databases (Interpro), allowing users to gain more insight into protein functions. The lower part of the output page (Fig 4B) shows the list of GO terms that are statistically enriched among the group of genome neighborhoods. Users can use this tool for, such as functional guidance, operon prediction, and protein evolution by observing the genome neighborhood patterns. The standalone version of this tool is freely available at https://github.com/asangphukieo/PhotoModGNN.

**Fig 4.**
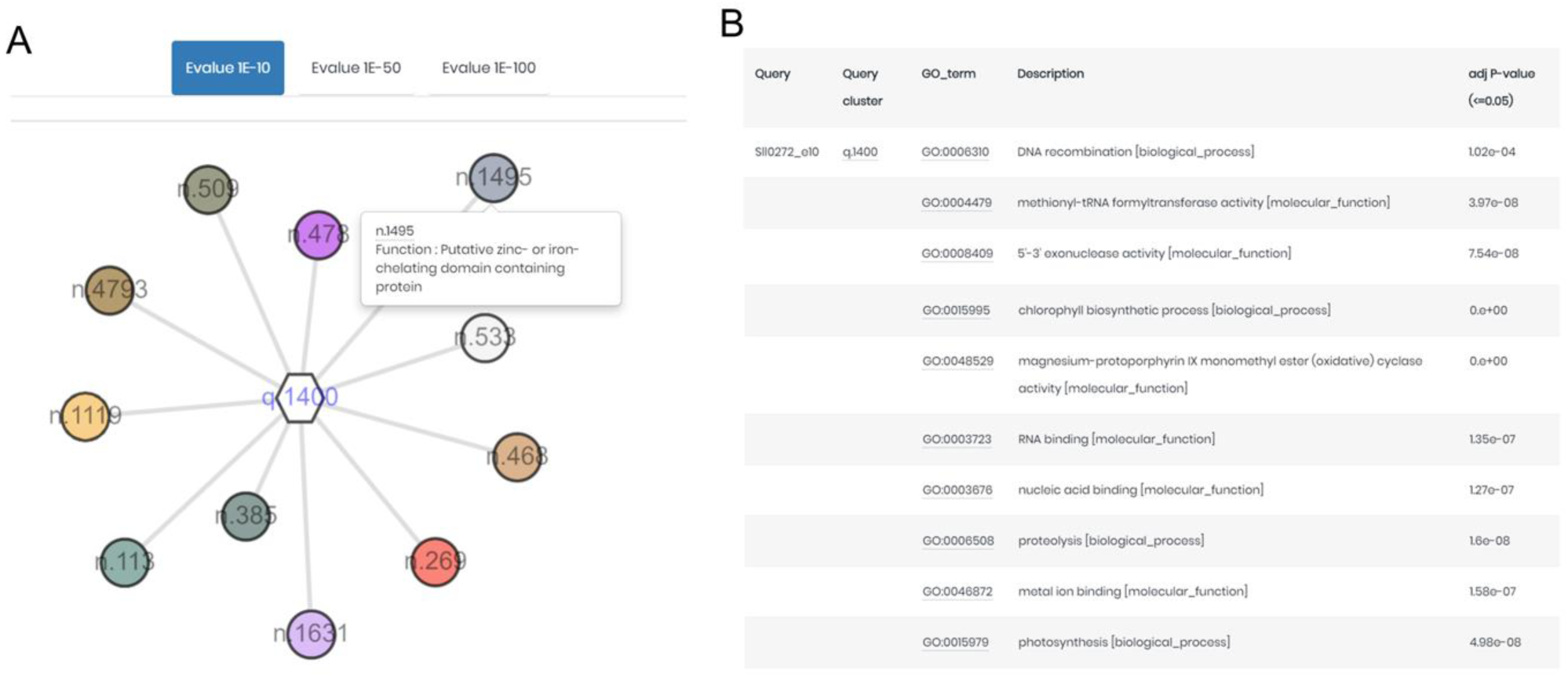
Example of GNN output from PhotoModPlus web server of sll0272. The GNN of sll0272 was shown in (A). The hexagonal node represents the sll0272 protein cluster, while the circular node represents the protein cluster from the neighboring gene. The size of the node is calculated according to the Phylo score (genome neighborhood conservation score), while the labeled number indicates a protein cluster ID. The GNN was displayed with an E-value of 1E-10. The list of GO terms that were enriched among the group of the genome neighborhoods was shown in (B).

We demonstrated the application of the GNN in Fig 4 by using the sll0272 protein sequence as an input. The GNN (Fig 4A) showed that sll0272 was conserved with 11 genome neighborhoods with moderate Phylo scores in range 2.37-3.96. The distribution of the Phylo scores collected from the entire dataset was shown in S7 Fig. The list of GO terms (Fig 4B) that were enriched among the group of the genome neighborhoods indicated that the query might involve in photosynthesis-related function and nucleic acid-related activity.

## Discussion

We demonstrated that the BLAST method performs efficiently when the sequences in the dataset are similar (≤70% sequence identity). However, if the sequences in the dataset are more diverse (≤50% sequence identity), the BLAST method suffers from lacking information resulting in low performance. Therefore, the machine-learning method plays an important role in this situation. We showed that the DeepGOPlus method, which combines the prediction based on sequence similarity and motif sequence-based neuron network model, performs better than the classical BLAST method in the diverse dataset. According to the fast, easy and efficient performance of the sequence-based approaches, they are suitable to use as a first tool to predict a protein function. When a more precise result is required, or no similar sequence is found in the database, it is necessary to use a different method. We can use the information from other sources to predict function, for example, protein-protein interaction, protein structure, and gene neighborhood [28]. However, gene neighborhood seems to be the cheapest and easiest data to find according to substantial reductions in the cost of genome sequencing. We developed PhotoModGO, which integrates protein clustering, genome neighborhood profile generation, and multi-label classification algorithm to classify photosynthesis subclasses. PhotoModGO outperformed the other two sequence-based methods in the diverse dataset, while the three methods showed comparable performance in the easy dataset. It indicates that PhotoModGO can assign the photosynthetic functions to proteins with high accuracy, even though the sequences of those proteins are highly diverse from training sequences. This situation is different from sequence-based approaches as DeepGO and BLAST perform very well in the easy dataset, but become worse in the diverse dataset.

Although we showed that PhotoModGO performs efficiently in the prediction of the photosynthetic subclass, it is important to mention about the limitations. Firstly, most of the photosynthetic-specific GO terms used in this study are in the biological process and cellular component categories (12 in biological process, 10 in a cellular component, and two in molecular function). We showed that PhotoModGO outperformed the sequence-based approaches, but we strongly believe that PhotoModGO performs worse on the molecular function category, in which the sequence-based methods generally work well [61]. Thus, in the PhotoModPlus web server, we also provide function prediction based on protein domain via interproscan as optional to improve the prediction quality. Secondly, gene neighborhood scoring is a time-consuming process. PhotoModGO calculates gene neighborhood conservation based on gene content-based phylogenetic tree, which was modified from Zheng, et al.’s study [62]. It requires the genome distance matrix based on shared gene content between genomes to calculate the score, therefore our dataset contains 154 genomes resulting in a huge amount of calculation (11,781 pairwise genome distance). Instead of using shared gene content, we might estimate the distance between genomes by using 16s rRNA, which can obtain from the public database such as the SILVA database (https://www.arb-silva.de), to increase calculation speed and to make the application easily expandable.

## Conclusion

According to the diversity of available data types and computational algorithms, many methods have been proposed for protein function prediction. Previously, PhotoMod was developed as an application for classifying photosynthetic proteins via the GNN feature. In this study, we created a new model, PhotoModGO, to predict photosynthetic function, including 24 subclasses. The new model employs a multi-label classification method and genome neighborhood feature for training. The model performance was shown to be better than sequence-based approaches. The web server platform, PhotoModPlus, was developed to increase accessibility to the applications, which consist of i) machine learning prediction to classify photosynthetic protein and its function, and ii) GNN construction to visualize genome neighborhood network to infer protein function.

## Implementation and availability

PhotoModGO is available as free software in Github at https://github.com/asangphukieo/PHOTOMOD2 and in Dockerhub at https://hub.docker.com/r/asangphukieo/photomod2. The PhotoModPlus web server is available at http://bicep.kmutt.ac.th/photomod.

## Acknowledgments

The authors thank Dr. Weerayuth Kittichotirat and Dr. Sawannee Sutheeworapong for insightful discussions and suggestions. We also thank Bioinformatics and Systems Biology Program and King Mongkut’s University of Technology Thonburi to support research materials and equipment.

## Funding

This work was partly supported by Petchra Pra Jom Klao Doctoral Scholarship (No: 13/2558) from King Mongkut’s University of Technology Thonburi and a research grant (NRMJ: 2559A30602134#60000108) from the National Research Council of Thailand (http://en.nrct.go.th). The funders had no role in study design, data collection and analysis, the decision to publish, and preparation of the manuscript.

## Competing interests

The authors declare that they have no competing interests.

## Supporting Information

**S1 Fig. Example of gene neighborhood and genome neighborhood network.** (A) Genes on the same strand were considered neighbors if they are within 250 bp intergenic distance or are overlapping. Additionally, the two clusters were merged into the same neighborhood gene cluster if they are in a range of 200-1000 bp in the divergent direction, based on the operon interaction concept [63]. (B) Gene neighborhoods are called from all of the genomes that contain a homolog of the query sequence. (C) After applying protein clustering and calculating the Phylo score, a genome neighborhood network (GNN) can be constructed. The hexagonal node represents the query, while the circular node represents its genome neighbors. The label in each node represents a protein cluster ID. The size of the genome neighbor node varies according to its Phylo score. There are three build-in protein clustering cutoffs (E-value: 1E-10, 1E-50, and 1E-100) available on the top of the network to define the level of homologous relationship of proteins.

**S2 Fig. Demonstration of nested 5×3 fold-cross validation.** The outer fold is shown in the upper part of the figure, while the nested fold is shown in the lower of the figure. For every outer fold, the training set (white color) is used to perform 3-fold cross-validation to find the best parameter set. The best parameter set is used to build a model again from the whole training set in the outer fold and tested with an independent test set (black color).

**S3 Table. List of 24 GO terms specific to the photosynthetic system.**

**S4 Table. P-value table of Wilcoxon signed-rank test in the performance comparison of multi-label classification of photosynthesis functions.**

**S5 Table. GO term prediction result of 17 novel photosynthetic genes from the DeepGoPlus model.**

**S6 Table. List of potential photosynthetic gene candidates predicted from PhotoModPlus web server.** The unknown protein-coding genes from Synechocystis sp. PCC 6803 genome (1,885 genes) were used as inputs in PhotoModPlus prediction. BLAST best match and E-value of 1E-25 were used as BLAST parameters. The potential candidates are selected from criteria i) predicted as a positive class with more than 80% probability of PhotoMod binary model and ii) containing at least one predicted GO term with more than 50% probability from PhotoModGO multi-label model.

**S7 Fig. Histogram of Phylo score distribution.** The Phylo scores were collected from the genome neighborhoods of photosynthetic protein dataset (A) and nonphotosynthetic dataset (B). Note that the zero value of the Phylo score indicates a nonconserved genome neighborhood.

## Notes

http://bicep.kmutt.ac.th/photomod

http://bicep2.kmutt.ac.th/photomod

### Competing Interest Statement

The authors have declared no competing interest.

